# High-glucose Induces Retinal Pigment Epithelium Mitochondrial Pathways of Apoptosis and Inhibits Mitophagy by Regulating ROS/PINK1/Parkin Signal Pathway

**DOI:** 10.1101/420653

**Authors:** Yuanping Zhang, Xiaoting Xi, Yan Mei, Xueying Zhao, Liqiong Zhou, Minjun Ma, Sili Liu, Xu Zha, Yanni Yang

## Abstract

Diabetic retinopathy (DR) caused visual performance degradation seriously endangers human beings’ health, uncovering the underlying mechanism might shed light on the discovery of DR therapeutic treatments. In this study, we found that the effects of glucose on retinal pigment epithelium (RPE) varies in a dose dependent manner, high-glucose promotes ROS generation and cell apoptosis, inhibits mitophagy as well as proliferative abilities, while low-glucose induces ROS production and cell mitophagy, but has little impacts on cell apoptosis and proliferation. Of note, the toxic effects of high-glucose on RPE are alleviated by ROS scavengers and aggravated by autophagy inhibitor 3-methyladenine (3-MA) or mitophagy inhibitor cyclosporin A (CsA). High-glucose induced ROS generation is merely eliminated by ROS scavengers instead of mitophagy or autophagy inhibitor. We also proved that high-glucose inhibits cell proliferation and promotes cell apoptosis by regulating ROS mediated inhibition of mitophagy. In addition, mitophagy associated proteins PINK1 and Parkin are downregulated by high-glucose or hydrogen peroxide treatments, which are reversed by ROS scavengers. Of note, Knock-down of PINK1 decreases phospharylated Parkin instead of total Parkin levels in RPE. Intriguingly, high-glucose’s inhibiting effects on cell mitophagy as well as proliferation and its promoting effects on cell apoptosis are reversed by either PINK1 or Parkin overexpression. Therefore, we concluded that high-glucose promotes RPE apoptosis and inhibits cell proliferation as well as mitophagy by regulating oxidative stress mediated inactivation of ROS/PINK1/Parkin signal pathway.

## Introduction

Diabetes mellitus (DM) is a common disease caused by hyperglycemia [1] and its complications such as diabetic retinopathy (DR) seriously endangers human beings’ health [2], DR degrades human’s visual performance, but the mechanisms are still unclear [3]. The retinal pigment epithelium (RPE) are the main cells of the retina and play important roles in the physiopathology of DR [4], and RPE treated with glucose are commonly used as an in vitro model for DR research [5]. Glucose treatment has various toxic effects on RPE, such as inhibition of cell proliferation [6] and induction of cell apoptosis [7], but the detailed mechanisms are still unclear.

Mitophagy is the selective degradation of mitochondria by autophagy and is critical for cells to maintain mitochondrial quality under environmental stress [8], which helps to protect cells from apoptosis and promote cell survival [9]. Recent study showed that DR progression is closely related with mitophagy, which prevents cells from apoptosis and alleviates DR development [10]. In addition, high glucose has been reported to impair autophagy and mitophagy in Neuro-2a cells [11], and induce retinal pericyte apoptosis and death [12], but there are no reports on low glucose’s effects on cell mitophagy and apoptosis. Therefore, It is reasonable to speculate that high glucose mediated inhibition of mitophagy might promote RPE apoptosis and death, which eventually causes DR pathogenesis.

Reactive oxygen species (ROS) levels are significantly increased by high glucose on primary cultured hippocampal neurons [13], and glucose metabolism also affects cell autophagy during starvation by regulating ROS levels [14], Recent study also found that ROS produced by oxidative stresses regulates a series of biological processes including cell apoptosis and autophagy, and ROS generation has also been reported to be related to DR progression [15]. Hence high glucose induced ROS generation might be pivotal for regulation of RPE mitophagy and apoptosis.

PTEN-induced putative kinase protein 1 (PINK1) /Parkin pathway is critical for regulating mitophagy and mitochondrial damage, its activation promotes mitophagy and protects cells from death and apoptosis [16]. Previous studies showed that ROS induced by cadmium promoted autophagy and mitophagy by regulating PINK1/Parkin pathway in mice [17, 18]. In addition, PINK1/Parkin signal pathway has been proved to play an important role in high glucose mediated regulation of RPE autophagy and apoptosis [19]. Of note, the mechanisms of PINK1 activating Parkin are still unclear, recent study reported that PINK1 phosphorylates Parkin [20], which is still need to be elucidated. Besides, the role of PINK1/Parkin in high-glucose mediated regulation of RPE cell death, proliferation and mitophagy are still unclear.

## Materials and Methods

### Cell culture and cell proliferation

Human retinal pigment epithelium cell line ARPE-19 was purchased from the American Type Culture Collection (ATCC, #CRL-2302™). ARPE-19 cells were diluted into the density of lxl0^4^/ml and seeded into 96-well plates, cells were then cultured under the standard conditions (37°C, 5% C02)for 12h. Different doses of glucose (15uM, 30uM, 50uM and 70uM) were added into the wells (each assay has three repetitions) and co-cultured with cells for 24h, 48h, 72h and 96h respectively. Cells were then collected and stained with Trypan Blue and counted under optical microscope. IOUL of CCK-8 solution was added into each well and incubated the plate for 1-4h in the incubator following the manufacturer’s instructions of Cell Counting Kit-8 (MedChemExpress Co., Ltd, USA). Before reading the plate, the plate was gently mixed on an orbital shaker for 1 minute, the Optical Density (OD) values were detected by a Gemini EM microplate reader (Molecular Devices, USA) at the absorbance of 450nm. The OD values were used to evaluate cell proliferative abilities after being treated by different doses of PQ at various time points.

### Real-Time qPCR

After treating ARPE-19 cells with different doses of PQ, TRIzol kit (Invitrogen, USA) was used to extract RNA from ARPE-19 cells following the manufacture’s protocol. Reverse transcription PCR by iScript cDNA Synthesis Kit (Bio-rad, Hercules, CA, USA) and Real-Time quantitative PCR by HiScript II Q Select RT SuperMix (Vazyme, China) were used to reverse and quantify relative GAPDH, MELK, PLK1, PLK4, PRC1, AURKA, AUPKB, CENPE, CCNB2, CENPF, PINK1 and Parkin expressions, the primers were designed and synthesized by Sangon Biotech Co., Ltd (Shanghai, China), primers are listed in Table1. Relative mRNA expression levels of genes were normalized by GADPH.

**Table 1.**
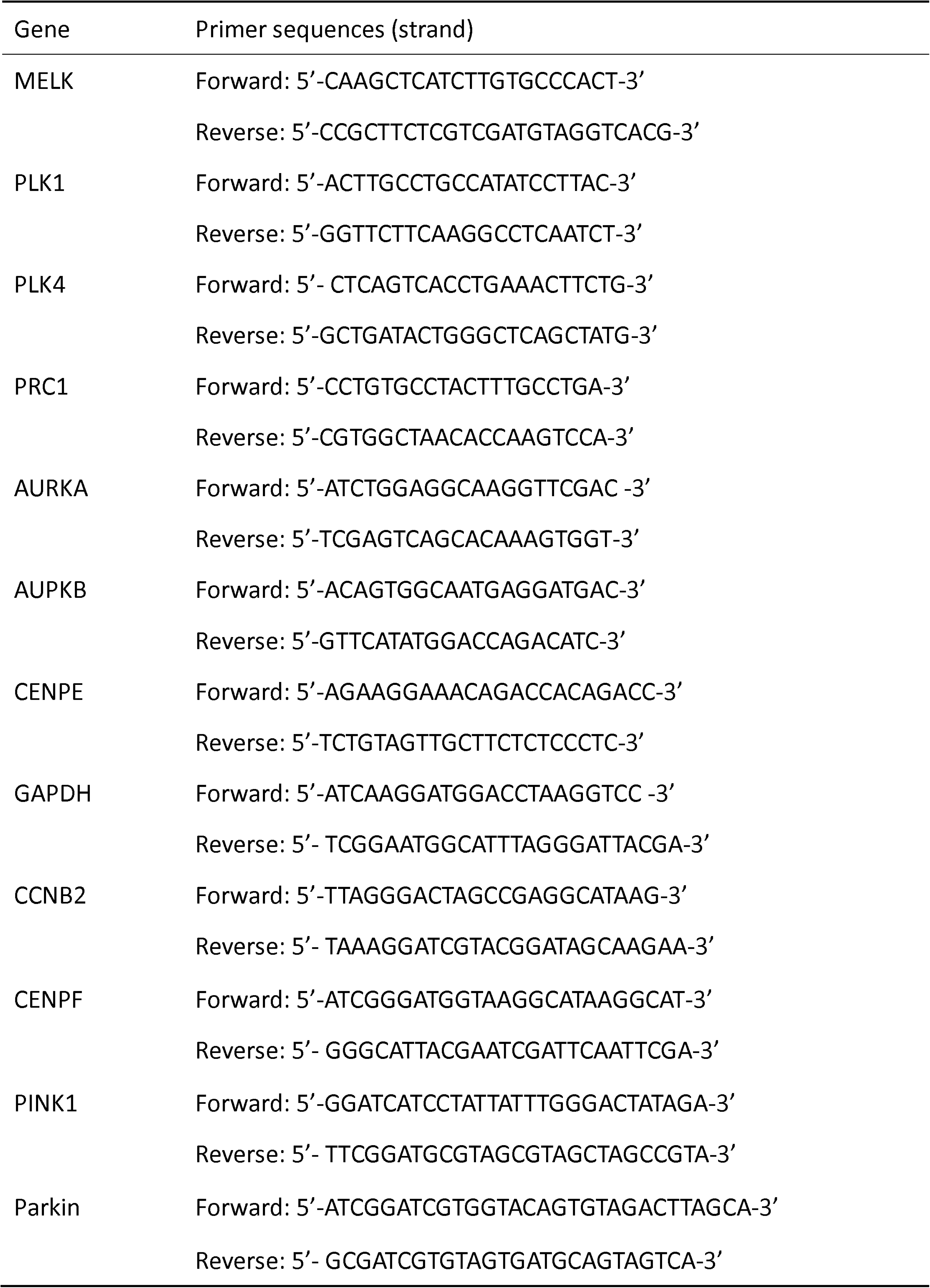
Quantitative PCR primers used in the study.

### Western Blot

The samples of ARPE-19 cells were collected, total protein was extracted using RIPA lysis buffer (Beyotime Biotechnology, Shanghai, China) and the protein concentration was determined by BCA Protein Assay Kit (Beyotime Biotechnology, Shanghai, China). Protein was then solubilized in 2x sample buffer, SDS-polyacrylamide gel electrophoresis was then performed to separate the targeted proteins. The proteins were then transferred to polyvinylidene difluoride (PVDF) membranes (Bio-Rad, Hercules, USA) and the membranes were incubated with 5% bovine serum albumin (BSA) for 60min at the room temperature. The primary antibodies of Anti-Cyt C (#SAB4500579, Sigma, USA), Anti-p62 (#P0068, Sigma, USA), Anti-CoxIV (#SAB 1405644, Sigma, USA), Anti-p21(#SAB4500065, Sigma, USA), Anti-Cyclin A2(#C4710, Sigma, USA), Anti-Cyclin D1(#C7464, Sigma, USA), Anti-Bax(#B8429, Sigma, USA), Anti-Bcl-2(#B3170, Sigma, USA), Anti-Caspase 3(#C8487, Sigma, USA), Anti-Caspase 9(#C7729, Sigma, USA), Anti-Caspase 8(#C4106, Sigma, USA), Anti-LC-3B (#8918, Sigma, USA), Anti-PINK1 (#P0051, Sigma, USA), Anti-Parkin (#P6248, Sigma, USA) and Anti-β-actin (#A5441, Sigma, USA) were employed to incubated with the membranes overnight at 4□. The anti-mouse IgG-peroxidase-conjugate secondary antibodies (Sigma, USA) were used to incubated with the membrane for 2 hours at the room temperature to combine to the primary antibodies. The bands were visualized by Enhanced Chemiluminescence Kit (Bio-Rad, USA) and ChemiDoc (Bio-Rad, USA), the expressions were quantified by Image J software. Each assay was repeated for at least 3 times to eliminate deviations.

### PINK1 and Parkin overexpression plasmids construction and transfection

To overexpress cellular PINK1 and Parkin expression levels, the coding sequence for the human PINK1 and Parkin genes were obtained from the total RNA of human HEK293T cells isolated by using TRIzolreragent (Takara Bio, Kusatsu, Japan) and then transcribed using iScript cDNA Synthesis Kit (Bio-Rad, Hercules, CA, USA). The sequences were them cloned into the pEGFP-N1 gene overexpression vectors, and the pEGFP-Nl-PINKl and pEGFP-N1-Parkin overexpression vectors were transfected into ARPE-19 cells by Lipofectamine 3000 transfection reagent (#13778150, Invitrogen, USA) according to the manufacturer’s instructions, the pEGFP-N1 empty vectors were used as the negative control. Western Blot was used to verify that the vectors were successfully transfected into the target cells.

### CRISPR-Cas9 technology was used to knockout PINK1 in ARPE-19 cells

CRISPR-Cas9 technology was conducted to knockout PINK1 gene according to the protocol from Dr. Zhang F. The two sgRNAs targeting human PINK1 were designed and constructed into pSgRNA (addgene#47108) using Bbs1 digestion (sgRNA sequences, F1: C AATTA AG G C ATG G A ATT CCC AT, Rl: AAACCTATTCCCAGAGTCAGTCA; F2: CATTCCTCCAGGACCCTAGGATGCAGT, R2: A A ACG TT CG A ACTCTG AC-GGTA). In addition, the constructed plasmids and pCas9-GFP (addgene#44719) were transfected into ARPE-19 cells by Lipofectamine 3000 transfection reagent (#13778150, Invitrogen, USA) according to the manufacturer’s instructions. Approximately lxlO^6^ cells were transfected with 1ug pCas9 plasmids, 1ug pSgRNAs and 0.2ug PKG-puro plasmid with puromycin resistance gene. Puromycin (1.5ug/ml) was then co-cultured with cells to select puromycin resistance cells. Total proteins were then extracted from the cells and Western Blot was used to verify the transfection efficiency of the plasmids.

### Flow cytometry

ARPE-19 cells treated with glucose were collected and prepared by GE Ficoll-Paque PLUS (GE, USA), cells were then treated with 10% DMSO and stored in -80□. Ex vivo cellular staining for Annexin-V and PI was implemented by incubating cells with specific dyes (ThermoFisher, USA) following the manufacturer’s instructions. Attune NxT Flow Cytometer (ThermoFisher, USA) was used to collect the data of cell necrosis, early apoptosis and late apoptosis. Each assay had at least 3 repetitions.

### Detection of ROS levels

ARPE-19 cells were treated with different doses of glucose, L-012 dye was used to detect extracellular NADPH oxidase-derived superoxide. In brief, ARPE-19 cells were diluted into approximately 4-6 x 10^4^ cells/well into 96-well plates (Thermo, USA) in phenol free DMEM medium (Sigma, USA) with L-012 at the concentration of 500uM according to our preliminary experiments (data not shown) for 10 min and luminescence was detected by a Gemini EM microplate reader (Molecular Devices, USA) at the excitation wavelength of 488nm and emission wavelength of 525nm respectively. Cellular ROS levels were next measured by Dihydroethidium (DHE) staining. Cells were washed with PBS twice and diluted, lOuM of DHE (Invitrogen, USA) was selected according to our preliminary experiments (data not shown) to incubate with the cells for 30min at 37°C without light exposure. After incubation, cells were washed with PBS and DM500 fluorescence microscope (Leica, Germany) was employed to observe ROS productions. The fluorescence intensity was quantified and calculated by Image J software.

### Statistical analysis

All the data collected in our experiments was showed as the mean ± standard deviation (SD), and the data was analyzed by SPSS 13.0 statistical software with one-way analysis of variance (ANOVA) for multiple groups and Student’s t-test for two groups. P<0.05 means statistical significance.

## Results

### Cell proliferation and ROS generation affected by different doses of glucose

Previous studies have reported that glucose induces ROS generation [14] and inhibits cell proliferation [21]. We first treated RPE with 15uM, 30uM, 50uM and 70uM glucose for 24h, 48h, 72h and 96h respectively, the cell counting and CCK-8 assay results showed that glucose inhibits RPE proliferation in a dose dependent manner (Figure 1A-D). After treating RPE with 15uM and 50uM glucose for 24h, Real-Time qPCR results showed that mRNA levels of the proliferation associated proteins (MELK, PLK1, PLK4, PRC1, AURKA, AUPKB, CENPE, CCNB2 and CENPF) are decreased by glucose in a dose dependent manner (Figure 1E). We next used Western Blot to detect cell cycle associated proteins (p21, Cyclin A2 and Cyclin D1) in RPE treated with 15uM and 50uM glucose for 24h, the results showed that low glucose (15uM) has little impacts on their levels, but high glucose (50uM) increases p21 and decreases Cyclin A2 as well as Cyclin D1 comparing to the control group (Figure 1F, G). RPE was then treated with 15uM and 50uM glucose for 4h, 8h, 12h and 24h respectively, ROS generation was detected by L-012 staining (Figure 1H, I) and DHE staining (Figure 1J, K), the results suggested that ROS levels are significantly increased by glucose in a dose dependent manner.

**Figure 1.**
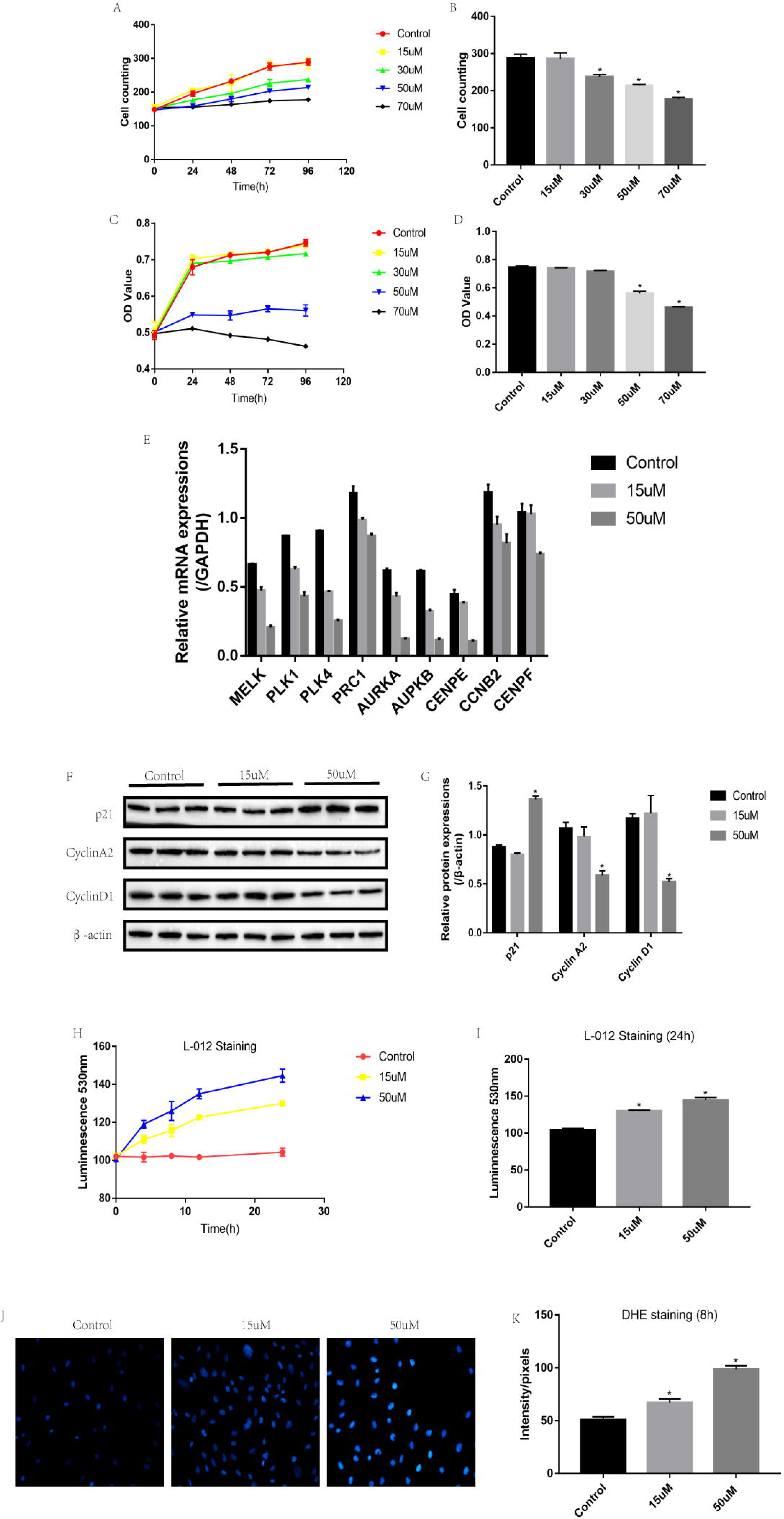
RPE proliferation and ROS generation influenced by different doses of glucose. (A) RPE treated with 15uM, 30uM, 50uM and 70uM glucose for Oh, 24h, 48h, 72h and 96h respectively, cell numbers were counted under inverted microscope. (B) RPE treated with 15uM, 30uM, 50uM and 70uM glucose for 96h, cell numbers were counted under inverted microscope. (C) RPE treated with 15uM, 30uM, 50uM and 70uM glucose for Oh, 24h, 48h, 72h and 96h respectively, CCK-8 assay was used to detect cell proliferation. (D) RPE treated with 15uM, 30uM, 50uM and 70uM glucose for 96h, CCK-8 assay was used to detect cell proliferation. (E) Real-Time qPCR was used to detect proliferation associated proteins, which were normalized by GAPDH. (F) After treating RPE with 15uM and 50uM glucose for 24h, Western Blot was used to detect p21, Cyclin A2 and Cyclin Dl, the bands represent protein levels. (G) Image J software was used to detect optical values of the protein bands and quantify the protein levels, which were normalized by P-actin. (H) RPE was subjected to 15uM and 50uM glucose for 4h, 8h, 12h and 24h respectively, and the superoxide release was monitored by chemiluminescence dye L-012. (I) L-012 staining was used to detect superoxide release in RPE treated with 15uM and 50uM glucose for 24h. (J) Images and (K) quantification of dihydroethidium (DHE) staining for intracellular ROS, the original objective magnification is 40X. Each assay in the experiments had at least 3 repetitions. (The data are presented as Mean ± Standard Deviation (SD), “*” means statistical significance)

### Cell mitophagy and mitochondrial pathways of apoptosis induced by glucose

Mitophagy is the selective degradation of mitochondria by autophagy, which is the protective mechanism for cells under environmental stress. After treating RPE with 15uM and 50uM glucose, the images photographed by electronic microscope showed that autophagosomes are induced by low glucose (15uM) instead of high glucose (50uM) comparing to the control group (Figure 2A). Western Blot was used to detect autophagy associated proteins, the results showed that LC3B-II/I ratio is significantly increased by 15uM glucose comparing to the control group, but LC3B-II/I ratio in 50uM group is significantly lower than 15uM group (Figure 2B, C). Low glucose (15uM) also decreases p62 in RPE, which is significantly increased by high glucose (50uM) (Figure 2B, D). To investigate whether LC3B is recruited to mitochondria, we successfully segragated cytosolic and mitochondrial sections in RPE treated with 15uM and 50uM glucose (Figure 2E). The results showed that LC3B-II/I ratio in the cytosolic section is increased by 15uM glucose and decreased by 50uM glucsoe comparing to the control group, which is in accordance with our previous results. However, LC3B is recruited to the mitochondrial section of RPE merely under the condition of RPE treated with 15uM glucose (Figure 2F). Low glucose (15uM) also has little impacts on RPE apoptosis ratio, which is significantly increased by high glucose (50uM) in a time dependent manner (Figure 2G,). Furthermore, high glucose (50uM) increases pro-apoptotic proteions including Bax, cleaved Caspase 9 and cleaved Caspase 3, while anti-apoptotic protein Bcl-2 is decreased by 50uM glucose comparing to the control group and 15uM group (Figure 2I, K, L, M, N, P). Intringuingly, pro-apoptotic protein caleaved Caspase 8 is not influced by either 15uM or 50uM glucose (Figure 2 J, N). Since Caspase 8 is involved in extrinsic apoptosis pathway [22], and Caspase 9/Caspase 3 axis participates in motochondrial apoptosis pathway [23], we next explored whether Cytochrosome C is released from mitochondria. Cytosolic and mitochondrial sections were successfully segragated from RPE treated with 15uM and 50uM glucose (Figure 2Q). Of note, we found that Cytochrosome C is merely released to the cytosolic section by treating RPE with 50uM glucose instead of 15uM glucose (Figure 2R).

**Figure 2.**
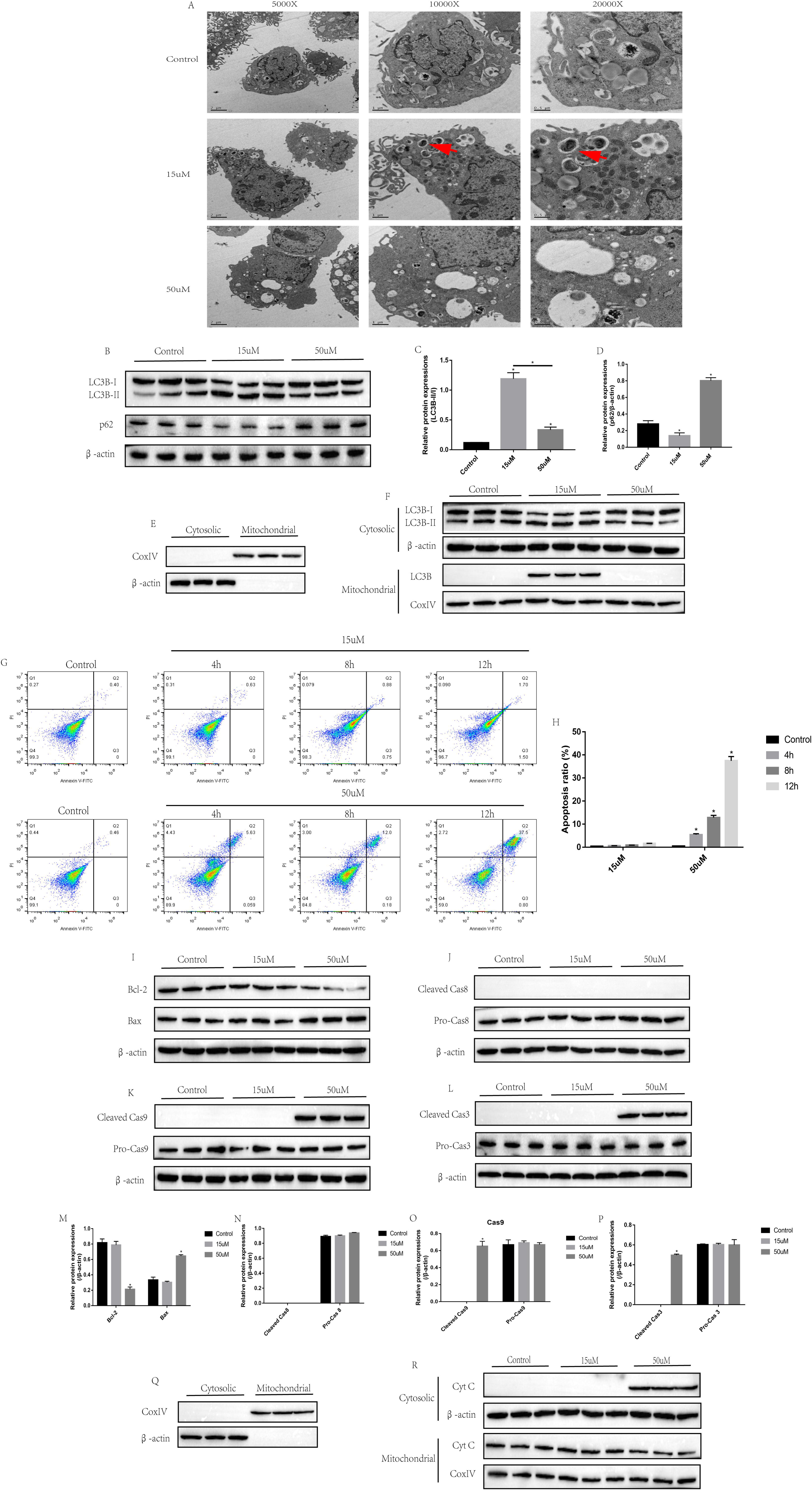
The effects of high- and low-glucose on cell mitophagy. (A) RPE was treted with 15uM and 50uM glucose for 4h, electronic microscope was used to observe cell autophagosomes in RPE. The red arrows indicate the autophagosomes. The images were 5000X, 10000X and 20000X respectively. (B) Western Blot was used to detect autophagy associated proteins (LC3B-II/I and p62) in RPE treated with 15uM and 50uM glucose for 4h, the bands represent protein levels. (C, D) Image J software was used to detect optical values of the protein bands and quantify the protein levels, which were normalized by p-actin. (E) CoxlV and p-actin were detected by Western Blot in cytosolic and mitochondrial of RPE respectively. (F) Cytosolic and mitochondrial LC3B was detected in RPE treated with 15uM and 50uM glucose by Western Blot. (G, H) RPE apoptosis ratio was detected by flow cytometry (FCM) after being treated with 15uM and 50uM glucose for 4h, 8h and 12h respectively. (I-L) Apoptosis associated proteins (Bcl-2, Bax, Caspase 8, Caspase 9 and Caspase 3) were detected by Western Blot, the bands represent relative protein levels. (M-P) The protein bands in (l-L) were quantified by Image J software, which were normalized by p-actin. (Q) CoxlV and p-actin were detected by Western Blot in cytosolic and mitochondrial of RPE respectively. (R) Cytosolic and mitochondrial cytochrome C (Cyt C) were detected by Western Blot, which were normalized by p-actin and CoxlV respectively. (The data are presented as Mean ± Standard Deviation (SD), “*” means statistical significance)

### Pretreatment with ROS scavengers and autophagy inhibitors under high glucose stress

Since oxidative stress is closely related with mitophagy [24], and mitophagy protects cells from apoptosis under environmental stress [25, 26]. We next explore whether high glucose inhibits RPE proliferation and promotes cell apoptosis by regulating oxidative stress mediated mitophagy. The L-012 staining results showed that ROS levels in RPE is significantly increased by high glucose, which is reversed by synergistically treating RPE with ROS scavengers NAC and ALC (Figure 3A). However, high glucose’s promoting effects on ROS generation are not influenced by eitehr autophagy inhibitor 3-MA or mitophagy inhibitor CsA (Figure 3B). Interestingly, we found that high glucose’s inhibiting effects on RPE proliferation are reversed by either NAC or ALA (Figure 3C), but aggravated by 3-MA or CsA (Figure 3D). In addition, high glucose promotes RPE apoptosis, which are reversed by either NAC or ALA (Figure 3E, F) and aggravated by 3-MA or CsA (Figure 3E, G). Furthermore, either NAC or ALA increases LC3B-II/I ratio (Figure 3H, I) and decreases p62 (Figure 3H, J) in RPE treated with high glucose. We next successfully segragated mitochondrial section in RPE (Figure 3K) and found that NAC promotes the recruitments of LC3B toward mitochondria (Figure 3L). Of note, we found that NAC’s alleviation on high glucose induced RPE apoptosis is abolished by synergistically treating RPE with 3-MA, on the other hand, 3-MA’s aggravation on high glucose induced RPE apoptosis is attenuated by synergistically treating RPE with NAC (Figure 3M, O). Similiarly, 3-MA or CsA’s aggravation on the inhibiting effects of high glucose on RPE is reversed by NAC, and vice versa (Figure 3N).

**Figure 3.**
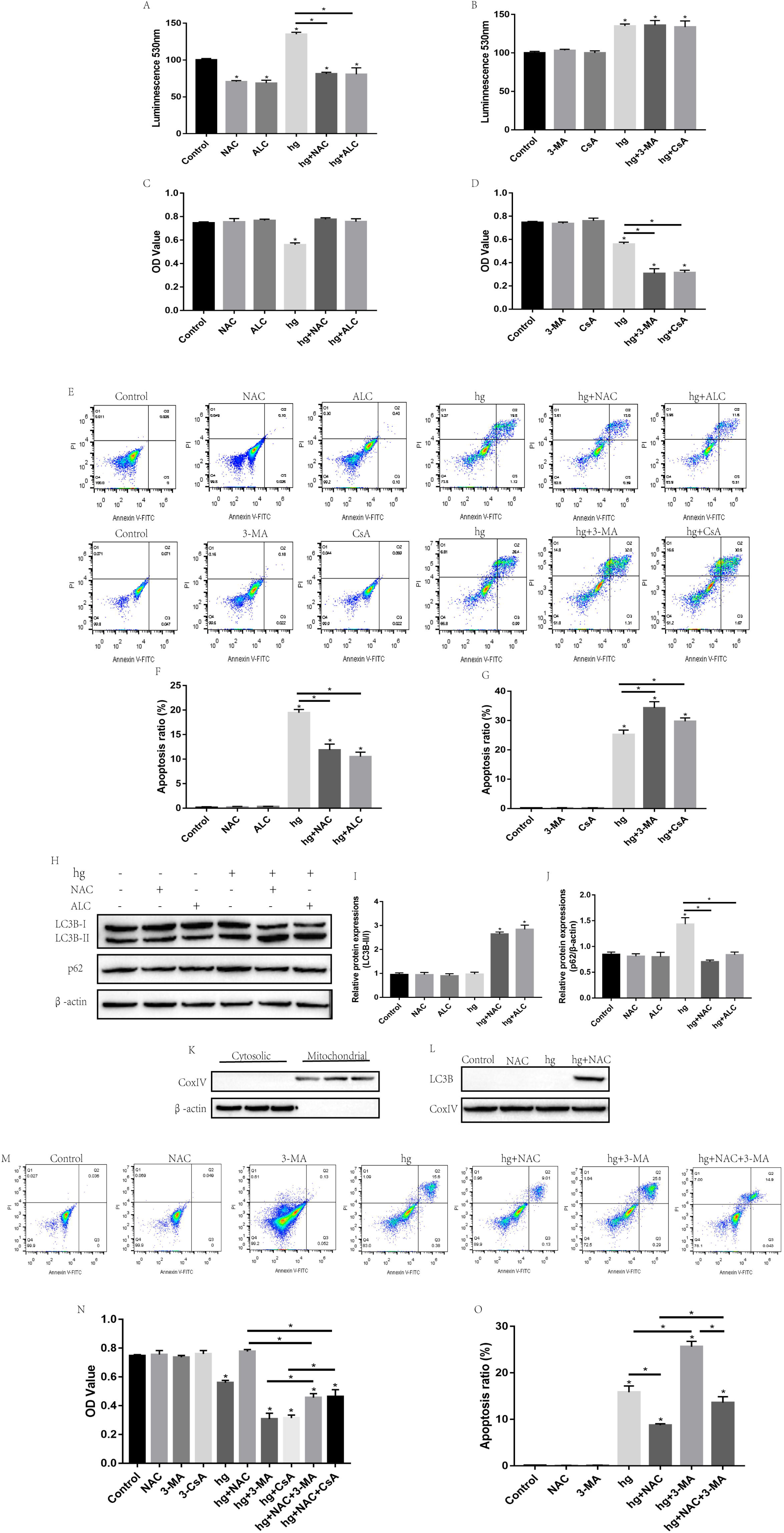
Pretreatment with ROS scavengers and mitophagy/autophagy inhibitors under high glucose stress. (A) RPE was subjected to 50uM glucose for 8h and synergistically pre-treated with ROS scavengers NAC and ALA, the superoxide release was monitored by chemiluminescence dye L-012. (B) RPE was subjected to 50uM glucose for 8h and synergistically pre-treated with autophagy inhibitor 3-MA and mitophagy inhibitor CsA, the superoxide release was monitored by chemiluminescence dye L-012. (C) RPE was subjected to 50uM glucose for 96h and synergistically pre-treated with ROS scavengers NAC and ALA, cell proliferation was detected by CCK-8 assay. (D) RPE was subjected to 50uM glucose for 96h and synergistically pre-treated with autophagy inhibitor 3-MA and mitophagy inhibitor CsA, cell proliferation was detected by CCK-8 assay. (E-G) RPE was subjected to 50uM glucose for 12h and synergistically pre-treated with ROS scavengers NAC as well as ALA, autophagy inhibitor 3-MA or mitophagy inhibitor CsA respectively, RPE apoptosis ratio was detected by FCM. (H) Autophagy associated proteins were detected by Western Blot, the bands represent protein levels. (I-J) The protein bands in (H) were quantified by Image J software, which were normalized by p-actin. (K) CoxlV and p-actin were detected by Western Blot in cytosolic and mitochondrial of RPE respectively. (L) Mitochondrial LC3B was detected by Western Blot, which was normalized by CoxlV (M, 0) RPE was synergistically treated with NAC and 3-MA, cell apoptosis ratio was detected by FCM. (N) PE was synergistically treated with NAC and 3-MA, cell proliferation was quantified by CCK-8 assay. (The data are presented as Mean ± Standard Deviation (SD), “*” means statistical significance)

### Mitophagy associated ROS/PINKl/Parkin signal pathway under high-glucose or hydrogen peroxide stress

PINKl/Parkin signal pathway has been reported to be involved in cell mitophagy [27] and is activated in RPE under high glucose treatment [19]. We next explored the underlying mechanisms of high glucose mediated activation of PINKl/Parkin signal pathway. Our Real-Time qPCR results showed that either high glucose or hydrogen peroxide inhibits PINK1 and Parkin mRNA levels in RPE, which are reversed by NAC or ALA (Figure 4A, B). The Western Blot results were similar with the Real-Time qPCR results, which indicated that high glucose or hydrogen peroxide inhibits PINK1 and Parkin in their protein levels, which are reversed by NAC or ALA (Figure 4C-H). Since PINK1 and Parkin showed a positive correlation, we next transfected RPE with PINK1 or Parkin overexpression vectors. The results showed that overexpressed PINK1 has no impacts on Parkin, and vice versa (Figure 41-K). Of note, PINK1 knock-out significantly decreses phosphorylated Parkin levels instead of total Parkin (Figure 4L-N), which is in accordance with the previous study [20]. In addition, either PINK1 or Parkin overexpression alone or high glucose has no impacts on LC3B-II/I ratio, but co-treating RPE with high glucose and PINK1 or Parkin overexpression vectors significantly increases LC3B-II/I ratio in RPE (Figure 4O, P). High glucose also induces p62 accumulation in RPE, which is reversed by either PINK1 or Parkin overexpression (Figure 4O, Q).

**Figure 4.**
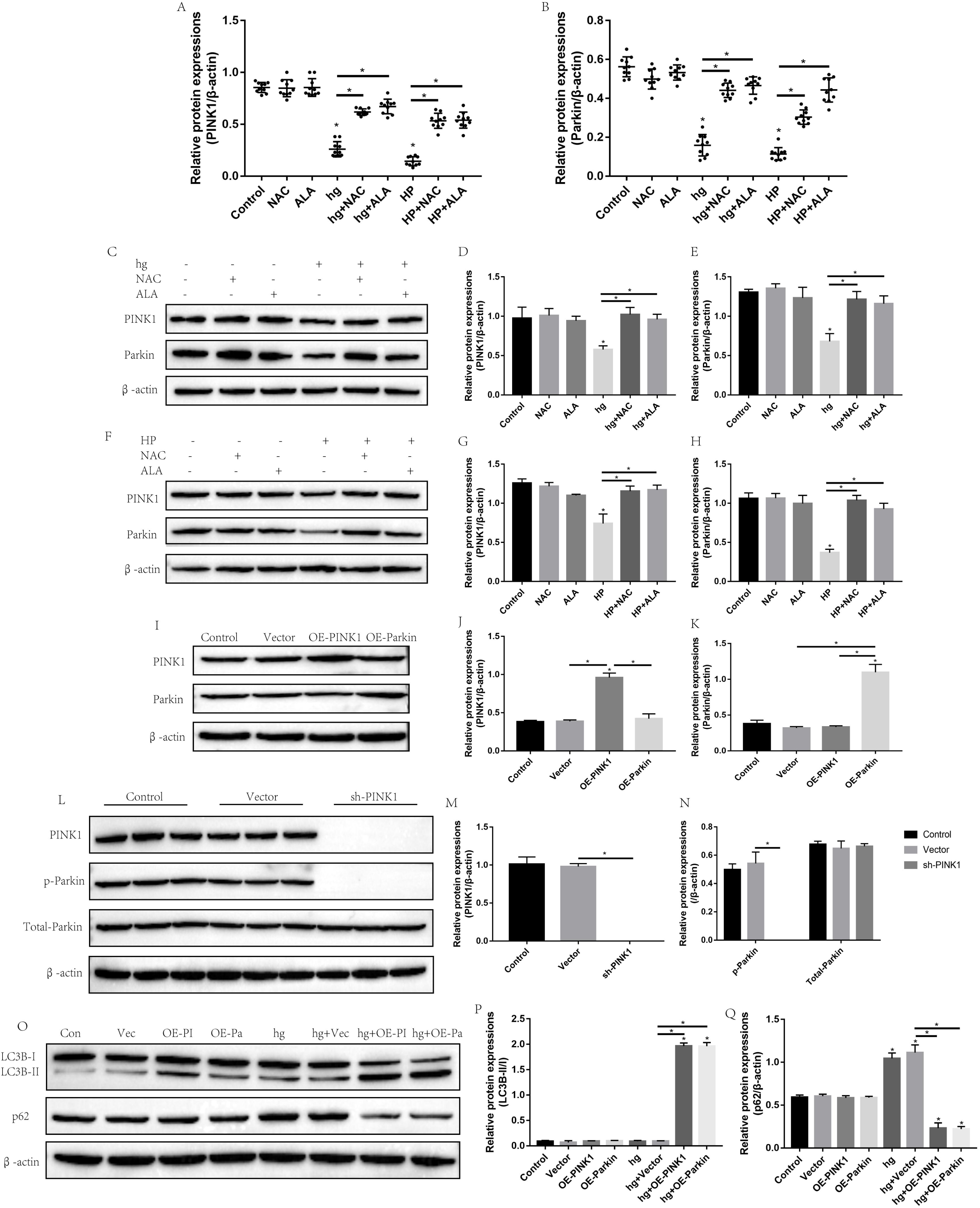
Involvement of mitophagy associated ROS/PINKl/Parkin signal pathway in RPE under high glucose or hydrogen peroxide stress. (A, B) The mRNA levels of PINK1 and Parkin in RPE treated with high glucose or hydrogen peroxidewere detected by Real-Time qPCR. (C) PINK1 and Parkin in RPE treated with high glucose and NAC/ALA were detected by Western Blot. (D, E) The protein bands in (C) were quantified by Image J software, which were normalized by p-actin. (F) PINK1 and Parkin in RPE treated with hydrogen peroxide and NAC/ALA were detected by Western Blot. (G, H) The protein bands in (F) were quantified by Image J software, which were normalized by p-actin.(I) PINK1 and Parkin in RPE transfected with PINK1 or Parkin overexpression vectors were detected by Western Blot. (J, K) The protein bands in (I) were quantified by Image J software, which were normalized by p-actin. (L) PINK1, total Parkin and phosphorylated Parkin in RPE transfected with PINK1 knock down vectors were detected by Western Blot. (M, N) The protein bands in (L) were quantified by Image J software, which were normalized by p-actin. (O) Autophagy associated proteins in RPE treated with high glucose and PINK1 or Parkin overexpression vectors were detected by Western Blot. (P, Q) The protein bands in (O) were quantified by Image J software, which were normalized by p-actin. (The data are presented as Mean ± Standard Deviation (SD), “*” means statistical significance)

### The role of PINKl/Parkin signal pathway in high glucose’s effects on RPE proliferation and apoptosis

We next investigated the involvement of PINKl/Parkin signal pathway in high glucose mediated regulation of RPE apoptosis and proliferation. The CCK-8 assay results showed that high glucose’s inhibiting effects on RPE proliferation are reversed by either PINK1 or Parkin overexpression (Figure 5A). Similiarly, our Western Blot results showed that high glucose promotes p21 and inhibits Cyclin A2 as well as Cyclin D1 in RPE comparing to the control group, which are significantly reversed by either PINK1 or Parkin overexpression (Figure 5B-E). FCM results showed that RPE apoptosis ratio is significantly increased by high glucose treatment, which is abrogated by either PINK1 or Parkin overexpression (Figure 5F, G). Furthermore, we detected mitochondrial apoptosis pathway associated proteins in RPE treated with high glucose and PINK1 or Parkin overexpression vectors. The results showed that co-treating RPE with high glucose and PINK1 or Parkin overexpression vectors decreases Bax, cleaved Caspase 9 as well as cleaved Caspase 3, and increases Bcl-2 comparing to the high glucose alone group (Figure H-L). In addition, we successfully segragated cytosolic and mitochondrial sections from RPE (Figure 5M), and we found that high glucose induced release of cytochrome C from mitochondria to cytoplasm is inhibited by either PINK1 or Parkin overexpression (Figure 5N-P).

**Figure 5.**
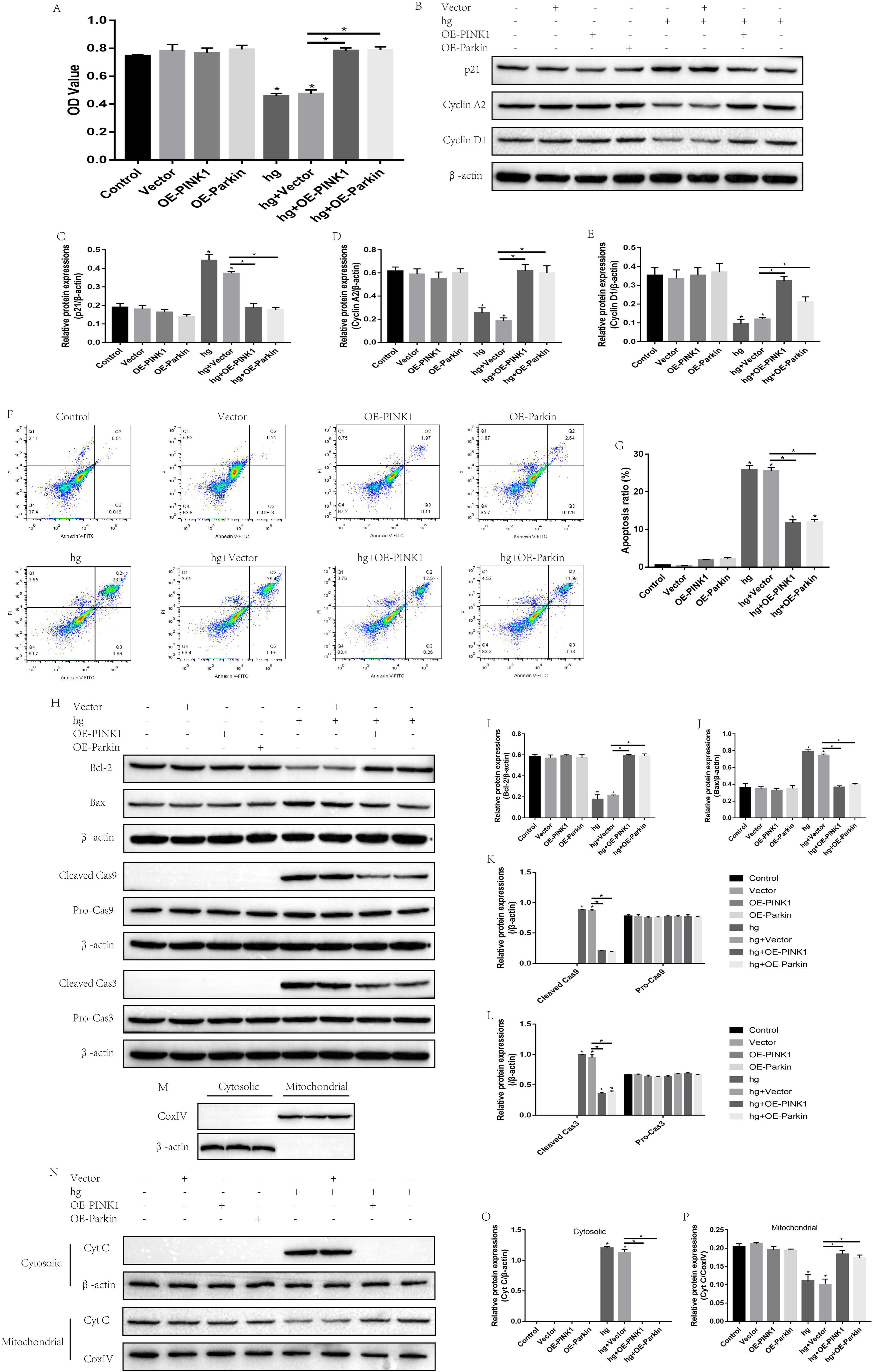
The role of PINKl/Parkin signal pathway in high glucose’s effects on RPE proliferation and apoptosis. (A) RPE was treated with high glucose and synergistically transfected with PINK1 or Parkin overexpression vectors for 96h, cell proliferation was detected by CCK-8 assay, the OD value represents cell proliferative ability. (B) Proliferation associated proteins of RPE in (A) were detected by Western Blot, the bands represent preotein expressions. (C-E) The protein bands in (B) were quantified by Image J software, which were normalized by p-actin. (F, G) RPE was treated with high glucose and synergistically transfected with PINK1 or Parkin overexpression vectors for 12h, cell apoptosis ratio was detected by FCM. (H) RPE was treated with high glucose and synergistically transfected with PINK1 or Parkin overexpression vectors for 12h, apoptosis associated proteins were detected by Western Blot. (I-L) The protein bands in (H) were quantified by Image J software, which were normalized by p-actin. (M) CoxlV and p-actin were detected by Western Blot in cytosolic or mitochondrial section of RPE respectively. (N) RPE was treated with high glucose and synergistically transfected with PINK1 or Parkin overexpression vectors for 12h, cytosolic and mitochondrial Cyt C were detected by Western Blot. (O, P) Proten bands in (N) were quantified by Image J software, which were normalized by p-actin or CoxlV respectively. (The data are presented as Mean ± Standard Deviation (SD), “*” means statistical significance)

## Discussion

Diabetic retinopathy (DR) seriously endangers human being’s health, and there are still no effective treatments for DR [28], uncovering the underlying mechanism might shed light on the discovery of therapeutic treatments for DR. RPE treated with glucose was commonly employed as the cell model for DR research [6, 29, 30]. It was reported that high glucose causes metabolic disorders of RPE by inducing oxidative stress [30]. Of note, the effects of glucose treatment on cell proliferation are different according to cell types, on the one hand, glucose promotes endometrial cancer cells [31] and mesangial cells [32] proliferation. On the other hand, high glucose induces rat glomerular mesangial cell injury [33]. However, the effects of glucose on RPE proliferation are still unclear.

Our results showed that RPE proliferation is inhibited by glucose treatment in a dose dependent manner (Figure 1A-D). Besides, mRNA levels of proliferation associated proteins are also decreased by glucose treatment in a dose dependent manner (Figure IE). The anti-proliferation protein p21 is increased and pro-proliferation proteins Cyclin A2 as well as Cyclin Dl are significantly decreased by high glucose instead of low glucose (Figure 1F-G). The results revealed that RPE proliferative ability is inhibited by glucose in a dose dependent manner. In addition, ROS generation is also induced by glucose in a dose dependent manner (Figure 1J-K), which is in accordance with the previous studies [34, 35].

We next investigated the effects of glucose on RPE autophagy and apoptosis. Our results proved that autophagy is merely induced by 15uM glucose instead of 50uM glucose (Figure 2A-D), which indicated that merely low glucose induces RPE autophagy. Of note, we found that LC3B is recruited to the mitochondrial section by treating RPE with 15uM glucose (Figure 2E-F), which suggested that low glucose specifically induces mitophagy. In addition, 50uM glucose significantly increases RPE apoptosis ratio (Figure 2G-H). Interestingly, high glucose increases cleaved Caspase 3 (Figure 2L, P) and Caspase 9 (Figure 2K, O), but has no impacts on cleaved Caspase 8 (Figure 2J, N). Further results also showed that 50uM glucose induces cytochrome C released from mitochondria to cytoplasm (Figure 2Q-R), which indicated that high glucose specifically induces mitochondrial patways of apoptosis in RPE.

Since oxidative stress was reported to be closely related with cell proliferation [36], apoptosis [37] and autophagy [38], we next investigated whether high glucose influences RPE proliferation, apoptosis and mitophagy by regulating oxidative stress. We found that ROS scavengers reverse high glucose’s effects on cell proliferation (Figure 3C), apoptosis (Figure 3E-F) and mitophagy (Figure 3H-L), which indicated that high glucose promotes cell apoptosis, inhibits cell autophagy and proliferation by increasing ROS levels in RPE. Notably, either autophagy inhibitor 3-MA or mitophagy inhibitor CsA aggravates high glucose’s inhibiting effects on cell proliferation (Figure 3D) and its promoting effects on cell apoptosis (Figure 3E, G), which indicated that autophagy or mitophagy play protective roles in high glucose induced RPE intoxication. Finally, we proved that high glucose inhibits RPE proliferation and promotes cell apoptosis by regulating oxidative stress mediated inhibition of cell mitophagy (Figure 3N, O).

Either high glucose or hydrogen peroxide is verified to decrease PINK1 and Parkin in RPE, which are significantly reversed by synergistically treating RPE with NAC or ALA (Figure 4A-H), which suggested that PINKl/Parkin signal pathway could be activated by high glucose mediated ROS generation. Surprisingly, we found that overexpressed PINK1 has no impacts on Parkin (Figure 4I-K), but knock-down of PINK1 specifically decreases phosphorylated Parkin instead of total Parkin in RPE cells, which suggested that Parkin is phosphorylated and activated by PINkl, our results are in line with the previous study [20]. In addition, we found that high glucose’s inhibiting effects on RPE mitophagy are reversed by either PINK1 or Parkin overexpression (Figure 4O-Q). Our results suggested that high glucose inhibits RPE mitophagy by regulating oxidative stress mediated inactivation of PINKl/Parkin signal pathway. Similiarly, we found that either PINK1 or Parkin overexpression reverses high glucose’s inhibiting effects on RPE proliferation (Figure 5A-E) and its promoting effects on cell apoptosis (Figure 5F-P).

## Conclusions

1. Low glucose induces RPE mitophagy and has little impacts on ROS generation, cell apoptosis and proliferation.
2. High glucose induces RPE apoptosis and inhibits cell proliferation as well as mitophagy by regulating oxidative stress mediated inactivation of PINKl/Parkin signal pathway.

## Acknowledgments

This study was supported by research grant 81860173 from the National Natural Science Foundation of China, and research grant 2018JS217 from The scientific research foundation of the Education Department of Yunnan Province.

